# *AMF-SporeChip* provides new insights into arbuscular mycorrhizal fungal pre-symbiotic hyphal growth dynamics at the cellular level

**DOI:** 10.1101/2023.06.29.546436

**Authors:** Felix Richter, Maryline Calonne-Salmon, Marcel G. A. van der Heijden, Stéphane Declerck, Claire E. Stanley

## Abstract

- Arbuscular mycorrhizal fungi (AMF) form symbiotic associations with the majority of land plants and deliver a wide range of soil-based ecosystem services. Due to their conspicuous belowground lifestyle in a dark environment surrounded by soil particles, much is still to be learned about the influence of environmental (i.e., physical) cues on spore germination, hyphal morphogenesis and hyphopodium formation in AMF.
- To fill existing gaps in AMF knowledge, we developed a new microfluidic platform – termed the *AMF-SporeChip* – to immobilise *Rhizophagus* and *Gigaspora* spores and confront pre-symbiotic hyphae with physical obstacles. In combination with timelapse microscopy, the fungi could be examined at the cellular level and in real-time.
- The *AMF-SporeChip* allowed us to acquire movies with unprecedented visual clarity and therefore identify various exploration strategies of AMF pre-symbiotic hyphae. We witnessed anastomosis formation involving directed hyphal growth in a “stop-and-go” manner, yielding visual evidence of pre-anastomosis signalling and decision-making. Remarkably, we also revealed a so-far undescribed reversible cytoplasmic retraction as part of a highly dynamic space navigation.
- Our findings demonstrated how AMF employ an intricate mechanism of space searching, involving reversible cytoplasmic retraction, branching and directional changes. In turn, the *AMF-SporeChip* is expected to open many future frontiers for AMF research.

## Introduction

Arbuscular mycorrhizal fungi (AMF) are a group of soil fungi belonging to the phylum Glomeromycota, forming a symbiosis with ca. 70-80% of all terrestrial plants. Laying the foundation for every soil-based ecosystem on earth (Bonfante *et al*., 2010; Cosme *et al*., 2018; van der Heijden *et al*., 2015), the fungus colonises root cells (i.e., the intraradical mycelium – IRM) and vastly extends into the soil (from 82 to 111 m cm^-3^ in prairie and 52 to 81 m cm^-3^ in ungrazed pasture (Miller *et al*., 1995)) to constitute the extraradical mycelium (ERM). This network of hyphae supplies the host plant with essential nutrients such as P and N, but also Zn, Cu and Fe (Smith *et al*., 2008), and receives between 4 and 20% of the total carbon synthesized by plants in return (Jakobsen *et al*., 1990; Shi *et al*., 2023). Furthermore, this immensely important mutualism increases the ecosystem’s overall resilience and facilitates plant productivity and plant diversity (van der Heijden *et al*., 2008). Moreover, AMF are not host specific and can interconnect different plant species forming so-called common mycorrhizal networks (CMNs) (Simard *et al*., 2012)).

Despite their importance, much is still unknown about the influence of environmental cues on spore germination, hyphal morphogenesis and hyphopodium formation in AMF, for example, with inoculation of AMF spores in the field often failing (Prado-Tarango *et al*., 2021; Berruti *et al*., 2016; Verbruggen *et al*., 2013). This is primarily due to the fact that the study of AMF and other soil-dwellers at the cellular and sub-cellular level proves to be challenging. The opacity of soil makes it impossible to observe these organisms in their natural habit and in real-time, rendering classic microscopy-based approaches either black-box experiments (e.g., end-point microscopy) or very simplified with limited resolution. The emergence, in recent decades, of *in vitro* cultivation techniques on root organs or whole plants has permitted to increase our knowledge on e.g., hyphal growth dynamics, three-dimensional architecture, anastomosis formation and hyphal healing mechanisms. The high numbers and density of hyphae, however, makes it more complex to resolve isolated events and a new development is required to investigate the behaviour of AMF at the single-hyphal level (e.g., hyphae morphogenesis, surface interactions with bacteria). Therefore, a new microfluidic tool was developed to study AMF in a visual, spatially-and temporally-resolved manner.

Microfluidic technology development for fungal research is a considerably young field, especially for studying hyphal behaviour and dynamics (reviewed in (Richter *et al*., 2022)). Being the first to explore this path, Nicolau and colleagues set out to study hyphal exploration strategies in micro-mazes, particularly with the filamentous model fungus *Neurospora crassa* (Held *et al*., 2009; Held *et al*., 2010; Held *et al*., 2011a). In recent years, this was developed further to accommodate more complex studies on filamentous fungi. Thomson *et al*. and Puerner *et al*. (Thomson *et al*., 2015; Puerner *et al*., 2020) for instance, utilised the deformability of poly(dimethylsiloxane) (PDMS) structures to measure forces exerted by growing hyphae. Further, microfluidic platforms have been developed to investigate interactions with other biological agents, such as bacteria (Stanley *et al*., 2014; Stockli *et al*., 2019), nematodes (Schmieder *et al*., 2019), bacteriophages (Ghanem *et al*., 2019) as well as fungal-fungal interactions (Gimeno *et al*., 2021). However, only one single study involving AMF has been published, which focusses solely on spore sorting (Srisom *et al*., 2020).

The use of microfluidic technology also allows to design microenvironments that mimic certain aspects of the natural habitat. Mafla-Endara *et al*. (Mafla-Endara *et al*., 2021), for example, created a soil-like microcosm featuring different geometries, species from various kingdoms, as well as water, air and soil particles as a substrate. In the course of that study, they observed how fungal hyphae help other organisms populate air-filled pockets in soil, hypothesising a water film in the mycosphere to be responsible for the transport. Further, gradients or niches of chemicals can be implemented for hyphal chemotaxis studies (Baranger *et al*., 2021). Ranking from highest to lowest in terms of complexity, field experiments, pot cultures and *in vitro* systems are all important experimental platforms for AMF research. Crucially, however, the microfluidics approach fills a significant experimental “gap”, providing a system with a high controllability of the experimental process and precise imaging, which allows for tailored patterning of the physical, as well as chemical, environment.

Herein, we introduce a new microfluidic device, termed the *AMF-SporeChip*, that was specifically designed to accommodate spores of *Rhizophagus* species for the study of spore germination events and exploration dynamics of pre-symbiotic hyphae for the first time. The devices were manufactured in-house using the elastomeric polymer, PDMS, which allows AMF hyphae to be confined to a single monolayer. Different micro-structures were included within the device design to mimic natural obstacles in soil (e.g., mineral particles, roots etc.), enabling hyphal dynamics to be explored upon confrontation with physical obstacles. Owing to the material’s near-perfect optical transparency, precise live imaging microscopy could be performed in real-time. To validate the compatibility of the device with AMF, we compared the germination and hyphal growth dynamics of 3 different *Rhizophagus* strains, both on-chip and on-plate, as well as demonstrating different germination behaviours between these strains. The flexibility of the system was also highlighted by (i) presenting a slightly modified device that can accommodate another AMF species, namely *Gigaspora margarita*, (ii) introducing fluorescent dyes (FM4-64 and Calcofluor White) into the microchannels for staining hyphae and (iii) demonstrating the suitability of the *AMF-SporeChip* for monitoring anastomosis formation in *Rhizophagus*. As a key result, we present an interesting and so far undescribed insight concerning reversible cytoplasmic retraction within presymbiotic hyphae, which amongst hyphal branching and directional changes, we consider part of their space searching strategy.

## Materials and Methods

### Chicory root and AMF culture

Three AMF strains, i.e., *Rhizophagus irregularis* MUCL 43194, *Rhizophagus irregularis* MUCL 41833 and *Rhizophagus intraradices* MUCL 49410, were provided by the Glomeromycota *in vitro* collection (GINCO, Belgium). The fungi were maintained in bi-compartmented root organ cultures (ROC) on Ri T-DNA transformed roots of chicory (*Cichorium intybus* L.). The roots were provided by the Glomeromycota *in vitro* collection (GINCO, Belgium) and cultivated on Modified Strullu and Romand (MSR) medium, stored at 27 °C in the dark in an inverted position and subcultured monthly. Spores of *Gigaspora margarita* BEG 34 were produced in pot co-culture with *Plantago lanceolata* (Cranenbrouck *et al*., 2005; Declerck *et al*., 1998). For more information and media preparation see the Supplementary Information (SI, Methods).

### Microfluidic device fabrication

The device designs were drawn in AutoCAD 2022 (Autodesk) and checked for correct structuring using KLayout (Köfferlein, 2006). The design was then printed to create a mylar® film photolithography mask by Micro Lithography Services Ltd., UK. Using these photomasks, two-layered master moulds were manufactured in a clean room (detailed in SI Methods).

To facilitate removal of the PDMS layer from the master moulds when casting devices, the wafers were silanised, i.e. made more hydrophobic. Therefore, the masters were carefully cleaned with an air gun and placed into a desiccator. In the centre of the desiccator, a glass vial containing 100 µl chlorotrimethylsilane (redistilled ≥99 %, Sigma Aldrich, UK) was placed. Using a vacuum pump, the desiccator was evacuated for 1 min and then closed off. The master moulds were left to react with the chlorotrimethylsilane for 1 h.

Using the master moulds, microfluidic devices can be produced on demand. All of the following steps were conducted in a laminar flow hood. Firstly, PDMS is mixed in a 10:1 ratio of base to curing agent (Sylgard 184, Dow Corning, USA) and degassed for ca. 45 min under vacuum in a desiccator. The master mould was fitted into a frame, the PDMS poured onto it and cured overnight at 70 °C. The PDMS was then removed from the mould, cut into slabs consisting of 2 device designs and holes for the in-and outlets were punched using a cork borer (Ø = 3 mm; Syneo, USA). To remove monomer remnants and to disinfect, the PDMS slabs were washed in 0.5 M NaOH and then 70 % ethanol, rinsing with sterile water between washing steps, and dried at 70 °C for 1 h. Next, the PDMS slabs as well as glass-bottomed Petri dishes (Ø dish = 35 mm, Ø glass = 23 mm; Fluorodish, World Precision Instruments, Germany) were plasma activated using a Zepto plasma cleaner (Diener Electronic, Germany; vacuum pressure 0.75 mbar, power 50%, 1 min) and bonded together, resulting in each Petri dish holding 2 devices. Exploiting the transient hydrophilicity of PDMS right after the plasma treatment, devices were filled with liquid hyphal medium (see SI Methods). Before use, devices were re-sterilised under UV light (254 nm) for 30 min.

### Device inoculation

The spore material of *R. irregularis* MUCL 41833 and MUCL 43194, as well as *R. intraradices* MUCL 49410 was obtained directly from the above-described bi-compartmented cultures (see SI Methods, *AMF culture*). All subsequent steps were performed in a sterilised laminar flow hood, under a stereoscope (EZ4 D, Leica, Germany). Using sterilised dissection needles, spore-bearing hyphae were extracted from a culture plate that is at least 3 months old and transferred into a 90 mm Petri dish filled with liquid MSR(-) medium (see SI Methods). Sterile tattoo needles (03 tight liner, pre-soldered, Barber DTS, UK) were used to cut the mycelium into smaller pieces containing 1-5 spores. Then, 5-6 of these spore pieces were transferred into each inlet of a device using a pipette (Pipetman G, 200 µl, Gilson, USA) that was set to 21 µl (to avoid overfilling of the inlets). A syringe (Henke-Ject, Luer Lock, 5 ml, HenkeSassWolf, Germany) connected to AlteSil High Strength Tubing (Bore: 1.00 mm, Wall: 1.00 mm, total diameter: 3 mm; Altec Extrusions Limited, UK) with a blunt tip syringe needle (1.28 mm, Shintop, China) and filled with liquid MSR(-) medium was carefully plugged into the inlet and spores were manually flushed into the device until all spores were inside of the channel and/or no more spores were coming from the inlet.

Spores of *Gi. margarita* were picked up using a pipette set to 20 µl and transferred into the inlet of the device. As the spores are detectable with the naked eye, it is easy to check visibly that only a single spore is in the pipette tip. The spores were then flushed into the device using the same method previously described for spores of *Rhizophagus*.

For staining experiments with fluorescent dyes, calcofluor white (to a final concentration of 0.5 µM) and FM4-64 (to a final concentration of 5 µM) were added to autoclaved liquid MSR(-) medium, sterile filtered and used to fill the microchannels as described above.

### Image analysis and quantification

Time-lapse images (used to produce growth and germination videos) as well as time-point large images (used for measuring growth rates) were obtained with two inverted microscopes (Eclipse Ti-U and Eclipse Ti-2, Nikon; SI Methods). Microscopy images were processed and analysed with Image J (Fiji) (Schindelin *et al*., 2012). To measure hyphal growth rates, the segmented line and measurement tool were used. The measured lengths were grouped into 5 intervals; 0.1-0.4 mm, 0.5-1.0 mm, 1.1-2.0 mm, 2.1-3.0 mm and >3 mm, and represented as stacked column diagrams using Origin 2021 (OriginLab). To account for the variation in number of spores loaded into the devices, the results were normalised. Therefore, the number of germination sites (instead of the number of spores) was determined and the absolute value of counts divided by the number of germination sites.

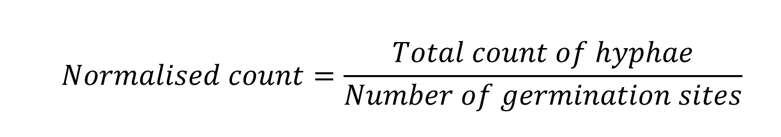

## Results

### Design and operation of the AMF-SporeChip

The *AMF-SporeChip* was designed to accommodate spores of different *Rhizophagus* strains, to enable high resolution dynamic imaging of spore germination and hyphal morphogenesis events at the cellular level. A two-dimensional (2D) sketch of the microchannel architecture is displayed in Figure **1a**, which features an inlet (Ø = 3 mm) connected to a “spore trapping region” (microchannel height = 100 µm) that then transitions into an “investigation zone” (microchannel height = 10 µm), with a total device length of 12.4 mm. Figure **1b** and **c** illustrate an overview of the *AMF-SporeChip*, where the change in microchannel height can be clearly evidenced via the addition of a fluorescein-containing solution, as well as how tubing is fitted into the device inlet for the introduction of AMF spores. The step change in microchannel height results in trapping of the AMF spores (Fig. **1d**), which usually have a diameter on the order of 10-100 µm for *R. irregularis* MUCL 41833, 20-160 µm for *R. irregularis* MUCL 43194 and 20-120 µm for *R. intraradices* MUCL 49410 (measured, Fig. **S1**), and importantly allows the much smaller, pre-symbiotic hyphae (Ø = 2-12 µm; measured, Fig. **S2**) to access (i.e., grow into) and be confined within the investigation zone in the z-direction. A microchannel height of 5 µm was also tested, with the intention of improving the confinement of hyphae further. However, the lower microchannel height made the channels more prone to collapse and the intended improvement in imaging was negligible. Therefore, a microchannel height of 10 µm was employed for the investigation zone.

**Figure 1:**
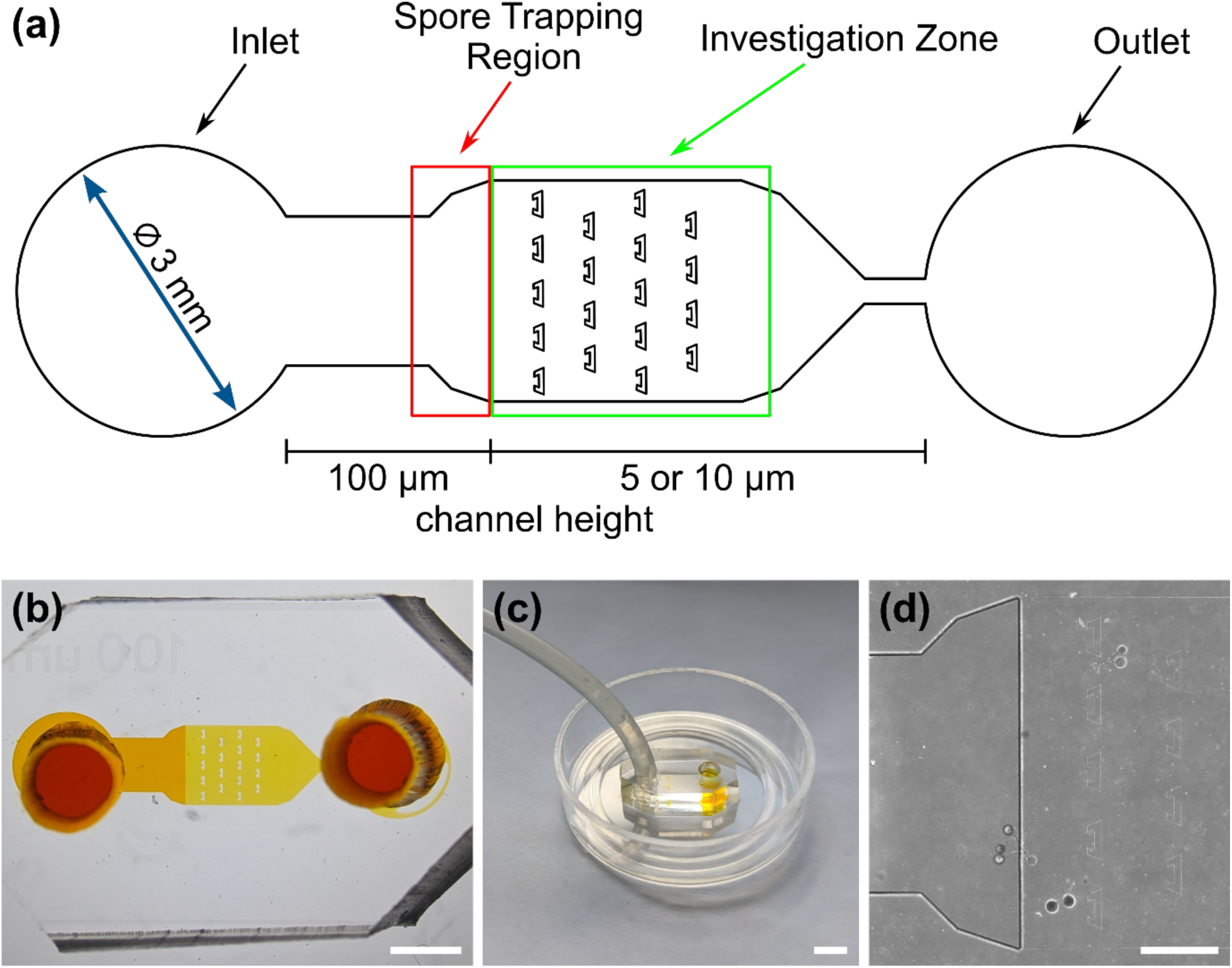
Design of the *AMF-SporeChip*. (a) Two-dimensional schematic of the microfluidic device showing the dimensions and structure of the spore trapping region and investigation zone containing micron-sized obstacles. (b) Real life image of the device filled with 66 mM fluorescein solution to highlight the channels and the change in channel height. (c) The *AMF-SporeChip* fitted with tubing at the device inlet for the introduction of spores. The outer diameter of the petri dish is 35 mm. (d) Brightfield image of the transition zone (microchannel height step-change) with trapped spores of *Rhizophagus irregularis* MUCL 41833. Scale bars: (b) 2 mm; (c) 5 mm and (d) 500 µm.

As PDMS is an elastomeric polymer and therefore flexible in nature, the microchannel widens slightly upon applying positive pressure when using a syringe to introduce spores into the *AMF-SporeChip* (Lee *et al*., 2012). Hence, smaller spores can be pushed into the investigation zone. Upon removing the applied pressure, the microchannel resumes its original state, thus trapping spores between top and bottom of the microchannel. For spores of *Gi. margarita* (Ø = 380-580 µm; measured, Fig. **S1**), the microchannel height of the inlet and spore trapping region was adjusted from 100 µm to ca. 350 µm. The aforementioned flexibility of the PDMS allows spores bigger than 350 µm to be accommodated within the trapping region of the microdevice, however not in the investigation zone.

The investigation zone contains an array of obstacles designed to provoke hyphal collision events (Fig. **1a**). A major advantage of the *AMF-SporeChip* is that the investigation zone can be modified in a bespoke manner to accommodate different types of physical obstacles that vary in their shape and size, or even possess lanes containing dead-ends or bottlenecks (Fig. **S3**). The obstacles varied in form, ranging from circular shapes that are either open (Pac-Man) or closed (circle), to rectangular-based (open-box) and triangular-based (restricted open-box) designs, thus offering increased or reduced chances for hyphae to “escape” from an obstacle. For measuring germination and growth dynamics, we utilised all designs equally to investigate how AMF hyphal tips react upon physical collision with their environment. In turn, the restricted open-box design proved to be most efficient in capturing hyphae. The array of interspaced obstacles provided an even inflow of medium and spore suspension, allowing for a balanced distribution of spores within the device, while giving plenty of opportunity for physical collisions.

### Monitoring spore germination and pre-symbiotic hyphal growth

To demonstrate the functionality of the *AMF-SporeChip*, a germination assay was conducted with (i) *R. irregularis* MUCL 41833, (ii) *R. irregularis* MUCL 43194 and (iii) *R. intraradices* MUCL 49410. Since visual representation and a detailed description of spore germination within these species are scarce, illustrative timelapse images of the germination process have been provided herein (Fig. 2, Movies **S1-3**), acquired from the examination of a total of 600 spores. When comparing the different strains, we observed quite distinct differences in germination and hyphal growth patterns as well as spore shape, despite being closely related. While *R. irregularis* MUCL 41833 and *R. intraradices* MUCL 49410 have a nearly perfect circular spore shape in our study, *R. irregularis* MUCL 43194 spores came in various, elongated shapes (Fig. 2; Fig. **S1**) Generally, it was observed that *R. irregularis* MUCL 41833 germinated with a single hyphal branch, as observed in Figure **2a**, with occasionally up to 4 branches emerging from the germination site. Subsequent hyphal growth was found to be relatively linear and straight. In contrast, *R. irregularis* MUCL 43194 spores tended to branch more frequently at or shortly after the site of germination (i.e., with typically 3-8 hyphae) as illustrated in Figure **2b**. Moreover, these pre-symbiotic hyphae possessed a curly and highly branched morphology. *R. intraradices* MUCL 49410 behaves similarly to *R. irregularis* MUCL 43194, with 3-6 hyphal branches emerging on average at or shortly after the germination site (Fig. **2c**). Another interesting observation included an active discharge of phase-bright cellular contents preceding germination (see Fig. **1b** and Movie **S2**). Occasionally, an internal collapse and emptying of the spore can be witnessed after a few days, after which the storage vesicles are ejected suddenly in one final load of cellular contents and nutrients into the attached hyphae (see Movie **S4**).

**Figure 2:**
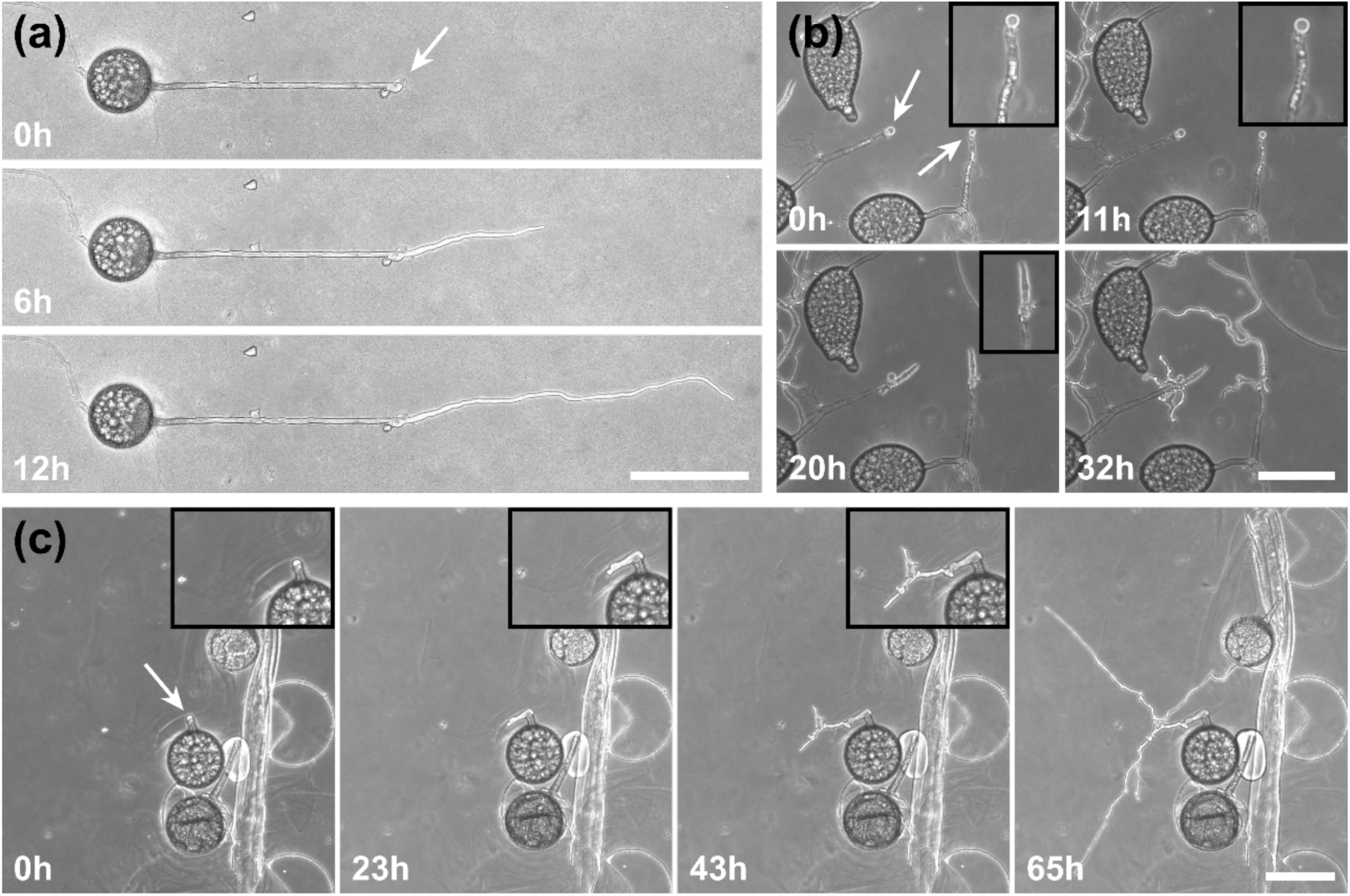
AMF germination on chip. Time-series illustrating the germination process of (a) *Rhizophagus irregularis* MUCL 41833, (b) *Rhizophagus irregularis* MUCL 43194 and (c) *Rhizophagus intraradices* MUCL 49410. White arrows indicate the germination site, with inset boxes in (b) and (c) illustrating an enlarged region of the germination event. Scale bars: 100 µm.

For a quantitative analysis of growth dynamics on-chip in comparison to on-plate, hyphae of all three species were imaged and measured as described in the Materials and Methods. The results are illustrated in Figure **3**. For *R. irregularis* MUCL 41833, the on-chip measurements indicated a constant increase in spore germination. This is illustrated by the orange-coloured bar segments, which represent freshly germinated and short hyphae with a length of 0.1-0.4 mm, as well as a similarly constant growth of hyphae to lengths of 0.5-1.0 mm (green) and few hyphae to lengths of 1.1-3.0 mm (purple and yellow) and longer (>3 mm, blue). For *R. irregularis* MUCL 43194, a greater increase in the number of hyphae was observed over time on-chip, reaching a plateau after day 4, with few hyphae then growing longer. *R. intraradices* MUCL 49410 showed a constant increase in the number of hyphae over 7 days, similar to *R. irregularis* MUCL 43194, however with barely any hyphae growing further than 1.0 mm and only a few having lengths between 0.5-1.0 mm.

**Figure 3:**
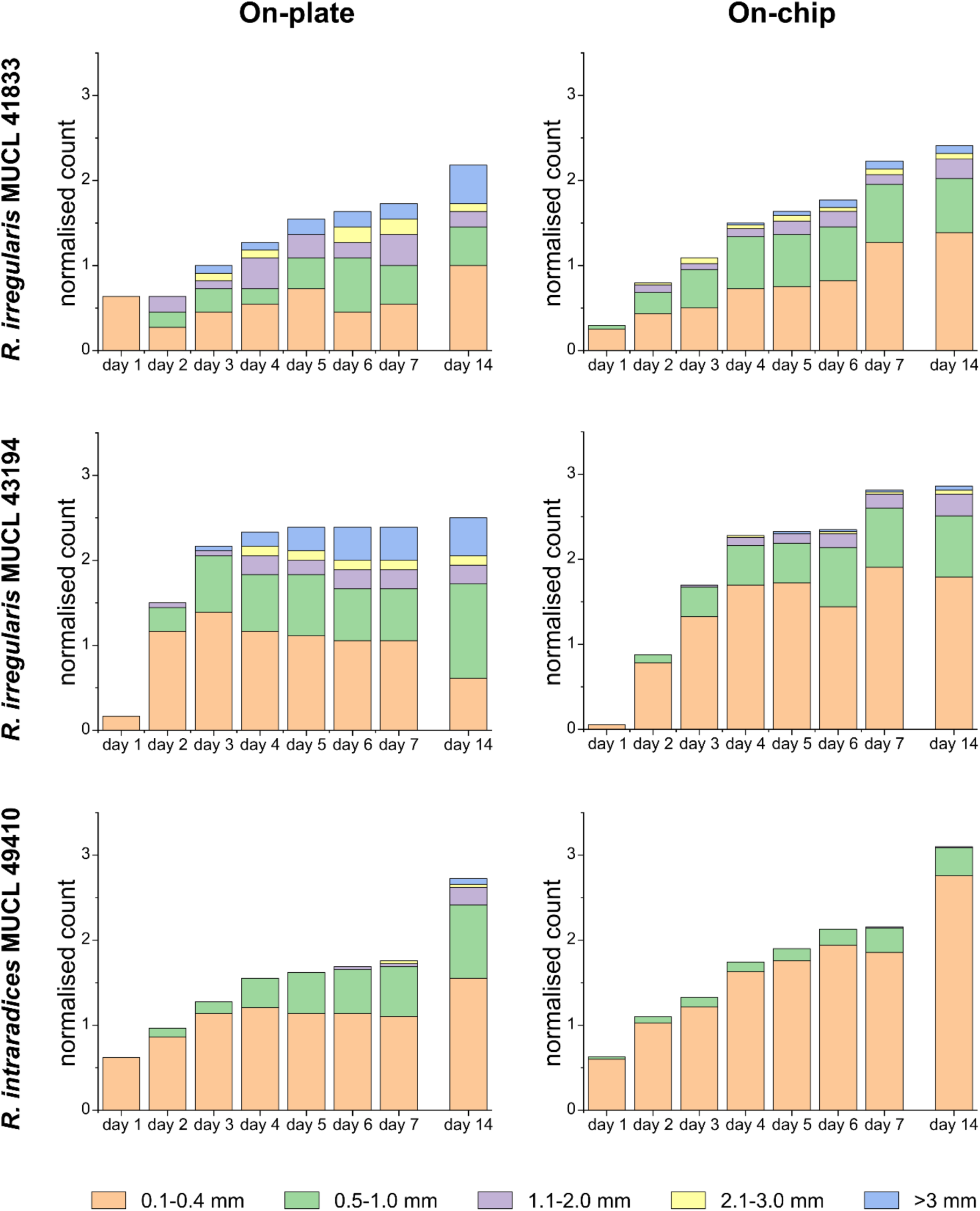
Hyphal growth assay on-chip and on-plate. Stacked column charts showing the growth behaviour of *Rhizophagus irregularis* MUCL 41833, *Rhizophagus irregularis* MUCL 43194 and *Rhizophagus intraradices* MUCL 49410 in the *AMF-SporeChip* and on MSR(-) culture plates for comparison. Hyphal lengths were measured using ImageJ, normalised (number of germination sites was determined, and the absolute value of counts divided by the number of germination sites) and categorised in 5 length sections. 6 devices with 40-60 germination sites in total per strain were used and 10-20 germination sites were found per plate.

Growth on-plate generally yielded longer, fast growing hyphae, reaching lengths of up to 20 mm, with slightly smaller total numbers of germination events, compared to on-chip. The trends illustrated by the stacked column charts, however, are very much comparable and similar between on-chip and on-plate assay. Between day 7 and day 14, there was little growth and few germination events observed on-chip for *R. irregularis* MUCL 41833 (ratio of normalised count for day 14/7 = 1.082), while on-plate showed both elongating hyphae and new germinations (ratio of normalised count for day 14/7 = 1.263). For *R. irregularis* MUCL 43194, there were barely any further germination events observed, both on-chip and on-plate (on-chip ratio of normalised count for day 14/7 = 1.017; on-plate ratio of normalised count for day 14/7 = 1.047), with no further hyphal growth on-chip but further elongation on-plate. With *R. intraradices* MUCL 49410, there were numerous fresh germination events observed after 7 days, for both on-chip and on-plate experiments (on-chip ratio of normalised count for day 14/7 = 1.437; on-plate ratio of normalised count for day 14/7 = 1.549). Few hyphae were found to elongate further after 7 days on-plate, but not on-chip.

To highlight the versatility of the device, we modified the *AMF-SporeChip* to accommodate spores of *Gi. margarita*. Compared to the model organism *Rhizophagus*, as well as other AMF, *Gi. margarita* differs mainly in its macroscopic spore size. Further, different anastomosis behaviour has been documented (de la Providencia *et al*., 2005). By increasing the height of the microchannel in the spore trapping region from 100 to 350 µm, *Gi. margarita* spores were trapped successfully with subsequent germination and pre-symbiotic hyphal growth observed within the microchannels (Fig. **4**). Upon encountering dead-end channels, the leading hypha bent and tracked the channel wall back in direction of the inlet, then branched in three different sites, ca. 350 µm, 720 µm and 1280 µm downstream from the hyphal tip. The hyphae then arrested growth and finally retracted the cytoplasm in a gradual manner, forming very pronounced septa (see Fig. **4** and Movie **S5**).

**Figure 4:**
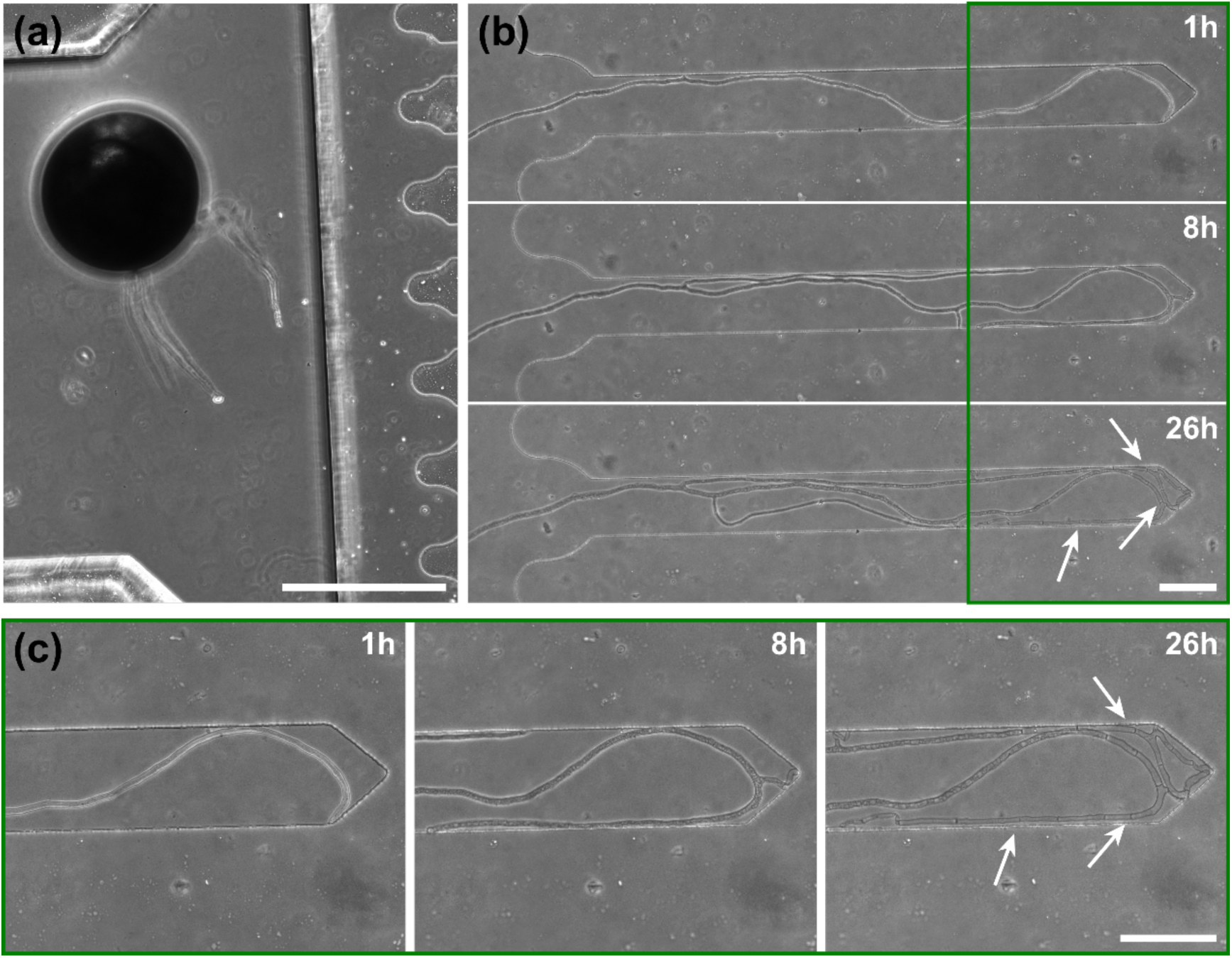
*Gigaspora margarita* growing in a microfluidic device. Shown are (a) a spore of *Gi. margarita* trapped in the inlet and (b) a timelapse series of a *Gi. margarita* hypha growing into dead-end channels. When the hypha hits the wall, it bends, branches off in several positions and starts retracting cytoplasm from trapped tips. White arrows show examples of very pronounced septa formed in the retraction process. (c) Magnifications of the dead-end of the microchannel. White arrows indicate examples of septa. Image (a) was taken 2 days before the start of the timelapse series in (b). Scale bars: (a) 500 µm, (b and c) 100 µm.

### “Stop-and-go” growth strategy prior to anastomosis formation

A highlight of the *AMF-SporeChip* involves the ability to observe fusion of hyphae to form a new cellular continuum, namely anastomosis. We witnessed both “tip-to-tip” and “tip-to-side” connections between hyphae originating from distinct spores, as well as from the same individuum. A representative depiction of anastomosis events observed within the *AMF-SporeChip* is shown in Figure **5**. The timelapse series presented show *R. irregularis* MUCL 43194 only; however, anastomosis events were also observed in *R. irregularis* MUCL 41833 and *R. intraradices* MUCL 49410. In Figure **5a** and **b** (and Movie **S6**), a tip-to-tip anastomosis event can be observed. The hyphae approach each other from opposing directions with occasional arrestation of growth and readjustment of the growth direction. Figure **5c** quantifies the growth of both hyphae over a 21 h period. The left hypha approached the other hypha in 4 main growth phases. Both duration, as well as distance covered, decreased from the beginning to the end of data recording, with a growth rate of ca. 0.3 µm/min for the first two growth phases and 0.1 µm/min for the latter two. The growth phases were interrupted by arrestation of growth for ca. 3 h, then ca. 4 h and before the final growth phase <2 h. Meanwhile, the right hypha barely grows at all, covering a total distance of ca. 12 µm in the 21 h time period. In spite of having plenty of space to explore, and thus avoid one other, they specifically target the other hypha, indicating a clear signalling and searching prior to anastomosing (Fig. **5c** and Movie **S6**).

**Figure 5:**
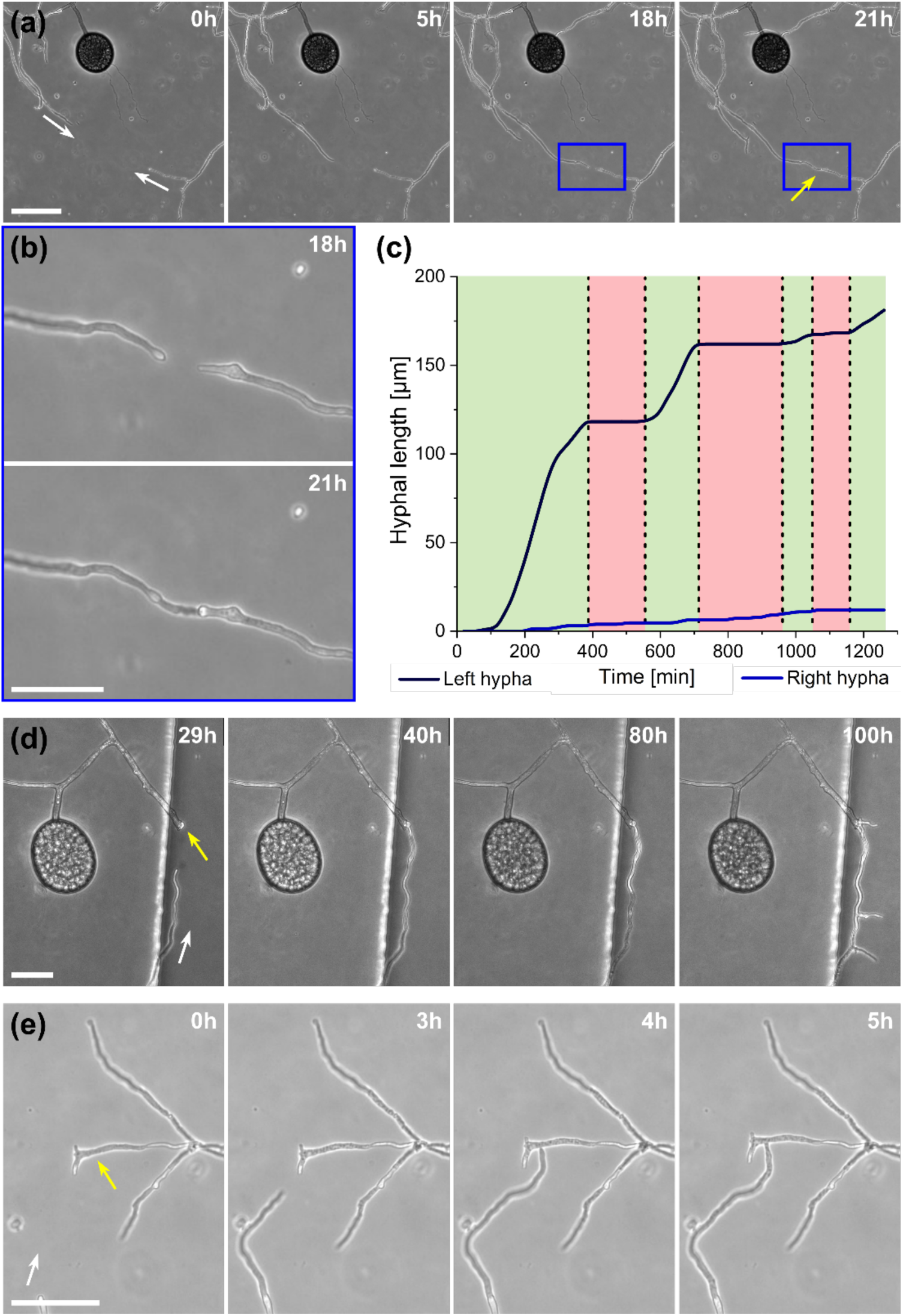
Anastomosis formation of *Rhizophagus irregularis* MUCL 43194 in microfluidic device. (a) Two hyphae approaching one other and forming a tip-to-tip anastomosis. (b) An enlarged image of the moment just before contact and upon contact in (a). (c) Graph highlighting the growth of both hyphae before fusing, which illustrates the “stop-and-go” growth strategy of the left hypha, with the hypha on the right-hand side barely growing at all. (d) Another example of a tip-to-tip anastomosis event where, however, the hypha coming from the bottom of the image attaches to an older hypha, which is just about to germinate. Further, it can be observed how they from a continuum, with cellular contents (phase-bright) being pushed into the bottom hypha from the direction of the spore, with subsequent branching. (e) A tip-to-side anastomosis. Yellow arrows indicate the anastomosis site, white arrows indicate the growth direction of anastomosing hyphae. Scale bars: (a), (d) and (e) 50 µm; (b) 25 µm.

In Figure **5d** (and Movie **S7**), a tip-to-tip anastomosis occurs in this instance, however, between a germination that is in progress and a hypha, rather than between two fully developed hyphae. The hypha approaching from the lower portion of the microscope image targets the end of an older hypha, which is just germinating, and fuses with the germination site. Following fusion, it can be clearly observed how a continuum is formed. Between timepoint 40 h and 80 h, cellular contents (phase-bright) are being pushed from the direction of the spore into the attached hypha. From this material, new branches are formed, concluding the process of unification. In Figure **5e** (and Movie **S8**), a tip-to-side anastomosis can be observed. A hypha approaches from the bottom left corner of the image and after 4 h attaches laterally to the other hypha. Almost immediately, the hyphae fuse and a cytoplasmic continuum is visible in the site of contact.

### Hypha behaviour upon encountering physical obstacles on the microscale

Inclusion of an array of obstacles, specifically the restricted open-box design, within the investigation zone of the *AMF-SporeChip* afforded new insights into AMF spore hyphal growth dynamics. Specifically, the following was observed upon collision of *R. irregularis* MUCL 41833 hyphae with these obstacles (see Fig. **6** and Movie **S9**). A hypha grew into the corner of an obstacle, i.e., a dead-end, resulting in the hypha bending slightly and arresting growth. It then retracted the cytoplasm from the trapped tip and subsequently formed a septum ca. 40 µm downstream from the hyphal tip. The septa was then breached, and cytoplasm pumped back into the trapped hyphal tip followed by a new branch emerging shortly behind the tip. This new branch then grew towards the opposite corner of the obstacle and proceeded to track the edge of the obstacle for a few hours, with its tip now pointed towards the spore it originated from. It then stopped growing and retracted its cytoplasm with at least 2 visible septa formed downstream. A new hyphal branch emerged from a position outside the obstacle, followed by a second branch growing from the same branching point but into the opposite direction a few hours later. Eventually the fungus stopped growing to the left-hand side (with respect to initial growth direction) soon after and retracted further cytoplasm from this and the original, trapped branch and continued growing towards the right-hand side (with respect to initial growth direction), forming a bifurcation around hour 57 after the start of the experiment.

**Figure 6:**
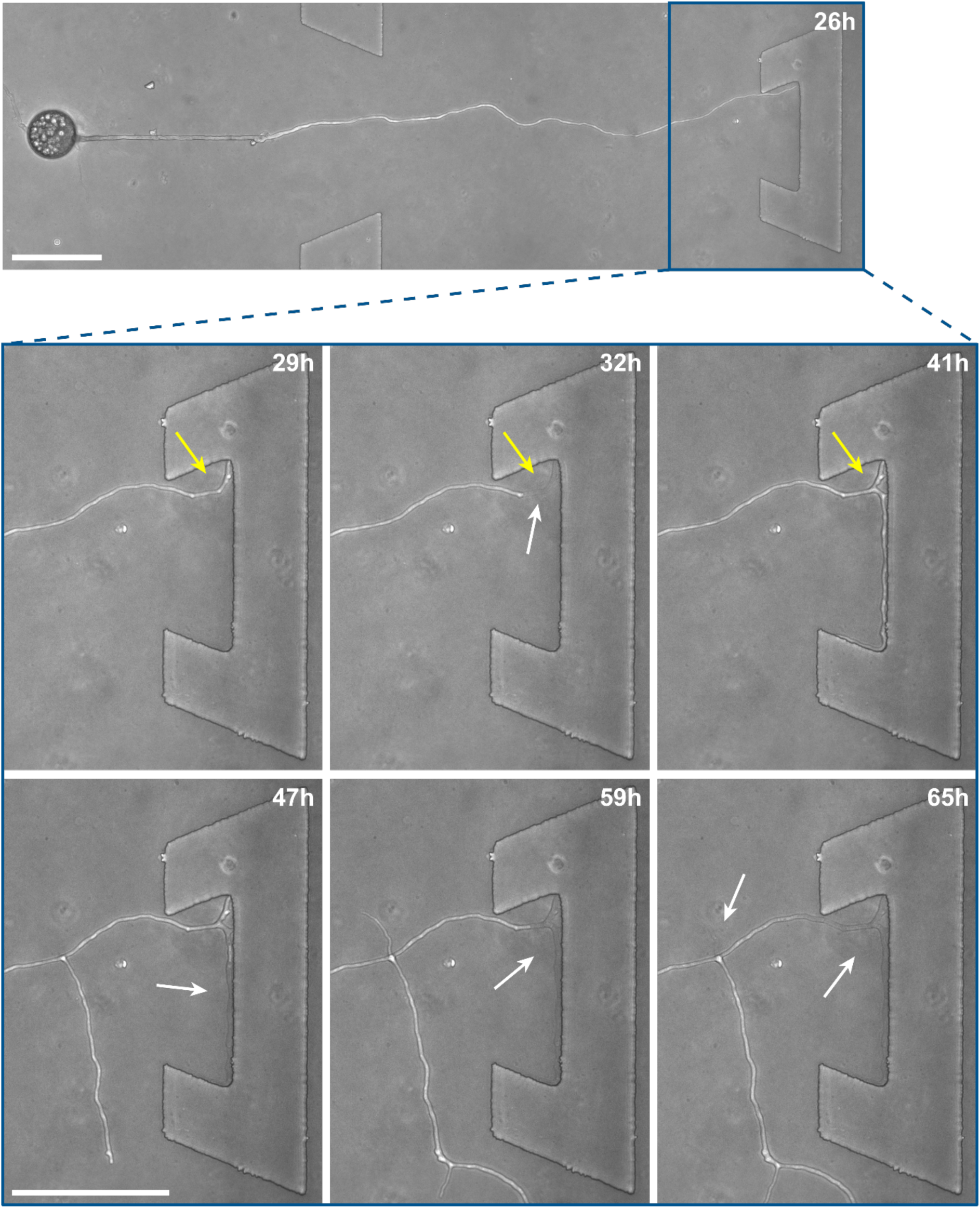
Hyphal collision event in the *AMF-SporeChip*. Here, a time-series of images illustrates the dynamic behaviour of a *Rhizophagus irregularis* MUCL 41833 hypha upon interaction with an obstacle in the microfluidic device. Timepoint 0 h was defined as the point of spore germination. The hypha grows into the obstacle and becomes trapped, after which a series of dynamic, reversible cytoplasmic retraction and branching events occur in several positions at varying time points. Yellow arrows indicate the “reversible” cytoplasmic retraction, white arrows indicate “empty” hyphae following cytoplasmic retraction. Scale bars: 100 µm.

To further highlight certain sub-cellular structures and characterise the nature of the hyphae observed in the cytoplasmic retraction, we employed two different fluorescent dyes, namely calcofluor white (CFW) and FM4-64 for end point staining. In Figure **7a**, all hyphae are visible in the phase contrast microscopy image, regardless of their developmental stage, however with differences observable, i.e., “normal” phase-bright hyphae and “empty” phase-dark hyphae, the latter of which is akin to what is observed after cytoplasmic retraction. Staining with the chitin specific dye, calcofluor white, reveals all hyphae observed in the phase contrast image (Fig. **7b**). In Figure **7c** only some hyphae are visible; here, the dye FM4-64 is employed, which stains lipid bilayers, i.e., membranes and organelles. Thus, only viable hyphae that contain all cellular contents remain detectable. Cytoplasmic retraction is followed by sequential formation of septa, which separate the cytoplasm-filled portion of a hypha from the empty portion. These septa are visible in phase-dark hyphae in the phase contrast image and are particularly pronounced in the CFW staining.

**Figure 7:**
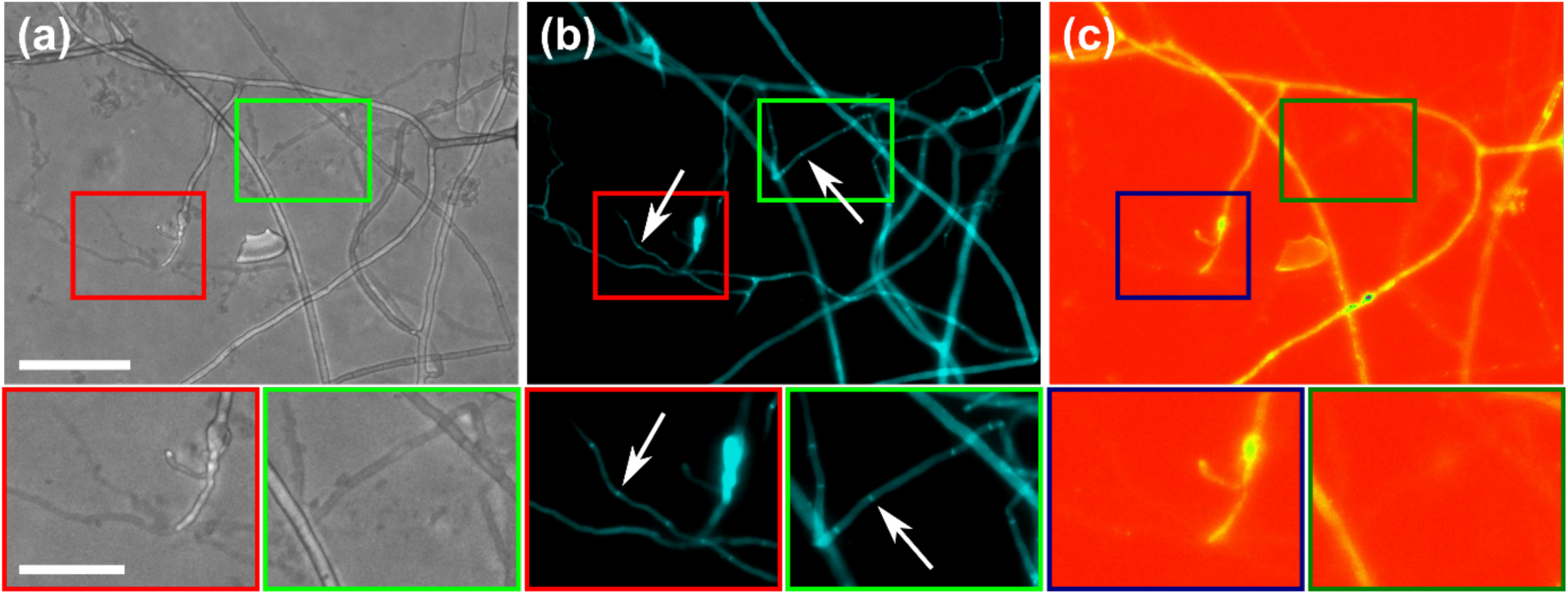
Differential staining of vital and empty hyphae. The images displayed refer to the same region of interest within the *AMF-SporeChip* and represent: (a) Phase contrast, (b) DAPI and (c) TRITC channels. The fluorescent dyes used in this study include calcofluor white (CFW), which stains chitin in the cell walls and FM4-64, staining lipid bilayers. White arrows indicate two exemplary hyphae which were emptied, with pronounced septa as artefacts of gradual retraction of cytoplasm. The images were edited using ImageJ/Fiji. The lookup tables “cyan” (b) and “spectrum” (c) were applied to improve the visibility of fluorescence intensity. Scale bar: 100 µm, 50 µm in zoomed-in images.

## Discussion

### The AMF-SporeChip provides high-resolution dynamic imaging of AMF in defined environments

Here, we present the first microfluidic device of its kind, namely the *AMF-SporeChip*, designed to accommodate AMF spores for studying germination and the dynamic behaviour of pre-symbiotic hyphal growth at the cellular level. The device was specifically designed to trap spores of *Rhizophagus* and later adapted for *Gi. margarita.* The trapping relied on two main aspects, the first being simply a difference in channel height, which kept bigger spores out of the investigation zone, while allowing the much smaller hyphae to grow in. Secondly, the high flexibility of PDMS allowed smaller spores within the population to be trapped at the entrance to the investigation zone. This trait of PDMS is actively utilised for cell trapping in microfluidics, mainly with yeast cells (Groisman *et al*., 2005; Lee *et al*., 2008; Zhang *et al*., 2012), but has also been used to measure force exerted by fungi-like oomycete hyphal tips (Sun *et al*., 2020). The application of microfluidic technologies for the study of yeast has received plenty of attention in the last two decades (Richter *et al*., 2022), yet for other types of fungi, this technique is still very new. Indeed, its utility has been demonstrated only in a handful of studies (Fukuda *et al*., 2021; Hopke *et al*., 2021; Held *et al*., 2011a), which focussed mainly on examining the space-searching behaviour of filamentous fungi or interactions between fungi and other microorganisms (Stanley *et al*., 2014; Schmieder *et al*., 2019; Gimeno *et al*., 2021), but never before with AMF.

Importantly, our devices featured obstacles within the investigation zone to mimic (micro) structures (i.e. soil microaggregates) in their natural habitat by blocking their growth path and provoking physical collisions. The various designs provided hyphae with differing opportunities to interact with or escape from an obstacle; the “restricted open-box” or the “dead-end lane” designs forced hyphae to arrest growth or change direction entirely. For AMF, this behaviour has never been studied before. Investigating how the fungal mycelium adapts to the physical conditions in soil is important for understanding a fungus’ lifestyle and eventually utilising this knowledge for potential applications in agriculture and horticulture/restoration as well as for the preservation of natural ecosystems.

### Microfluidics-assisted imaging yields detailed visuals of spore germination in real-time

Germination of AMF spores has been described in the literature (Souza, 2015), however, high-resolution visual material is scarcely available. To demonstrate the strength of the microfluidic approach and as proof-of-concept, we captured high resolution timelapse videos of spores of the three *Rhizophagus* strains studied, as well as with *Gi. margarita*.

Interestingly, distinct germination and branching patterns were observed between *R. irregularis* MUCL 41833 and 43194 (despite being the same species, and besides differences in shape and size), which suggest different exploration strategies. While *R. irregularis* MUCL 41833 germinated with 1-4 rather straight hyphae from the germination site, *R. irregularis* MUCL 43194 germinated with between 3-8 curly hyphae, which readily branched again shortly after the germination. Measuring hyphal elongation every 24 h for a week further revealed that *R. irregularis* MUCL 41833 has a growth pattern which is relatively balanced between new germination and hyphal elongation, whereas *R. irregularis* MUCL 43194 tended to germinate a lot more over the first 3 days and then resulted in elongation of some of these hyphae, corroborating the visually observed growth behaviour in both strains. It appears this strain first establishes a base around the spore and then starts exploring with single runner hyphae, while *R. irregularis* MUCL 41833 does not hesitate and starts exploring straight away with runner hyphae shooting randomly from the spore. The physiological stage (i.e., spore age) was not taken into consideration. For both strains, hyphal elongation proceeded after day 7 but rarely were any new germinations observed. *R. intraradices* MUCL 49410 behaves in a similar manner to *R. irregularis* MUCL 43194, with typically 3-6 curly and branched hyphae emerging from one germination site, which, however, barely ever extended beyond 1 mm. Fresh germination events on the other hand continued to occur numerously even beyond day 7 until the end of the experiment.

For all three strains studied, the on-plate results followed closely the findings from the on-chip experiments, with the exception of increased hyphal elongation on-plate. This difference in growth is a pre-described observation, which occurs in microfluidic devices and could be explained by two observations: (i) the microchannels are saturated with culture medium, i.e., the fungi are submersed in liquid, while on-plate they grow in an environment that has a solid phytagel base, with a liquid film on top as well as direct contact with air; (ii) the hyphae are physically confined inside microchannels, which has been suspected to influence hyphal growth (Baranger *et al*., 2020). Filamentous fungi are known to possess mechanosensory properties (Kumamoto, 2008) allowing them to sense the confinement and adapt their growth accordingly. This sensing happens both on a mechanical level, upon direct contact with the wall or the obstacle, as well as chemical level, where the fungus recognises the surface functionalisation (Held *et al*., 2011b). Overall, we can conclude from our results, however, that the germination and growth behaviour in the *Rhizophagus* strains is very much comparable between on-chip and on-plate and hence our microdevices are suitable to study these fungi. Further, the devices were modified to accommodate the much bigger spores of *Gi. margarita* and it was demonstrated that they, too, can germinate in the microdevice, illustrating the flexibility of our platform.

### Anastomosis formation involves directed growth with readjustments in “stop-and-go” manner

Another phenomenon implicating hyphal space searching strategies is anastomosis formation. Filamentous fungi are able to connect their hyphae with another individuum of the same strain or in a “self-self” manner (i.e., hyphae from the same individuum) (Novais *et al*., 2017) (Glass *et al*., 2000). Even interspecies anastomoses have been reported (Roca *et al*., 2004). Fungi anastomose to interconnect and expand their mycelial network for exploring and foraging, to exchange nutrients (Fleißner *et al*., 2008) and genetic material, (Friesen *et al*., 2006) or to heal damages in the hyphal network (de la Providencia *et al*., 2005). Here, anastomoses were observed between different individuals of the same strain as well as self-self, and formed in a tip-to-tip or tip-to-side manner. Our high-resolution imaging approach revealed how two hyphae hesitantly approached one other from opposing directions in a stop-and-go manner, occasionally halting to readjust growth direction. It is clear that these hyphae actively target each other since there is plenty of space for avoidance, which suggests an underlying signalling and decision-making that leads to tropism for one other. This was further corroborated by the occasional observation of hyphal contact that did not lead to successful anastomosis, indicated by cytoplasmic retraction and septation as earlier described by Giovannetti et al. (Giovannetti *et al*., 1999). In certain filamentous fungi, it has been reported that there are even specialised hyphae for fusion (McCabe *et al*., 1999; Read *et al*., 2009).

### Physical obstacles trigger dynamic hyphal reactions involving reversible cytoplasmic retraction

The obstacles featured in our devices were designed to trigger collision events, which we hypothesised would provoke a response in AMF akin to that described for other filamentous fungi (Aleklett *et al*., 2021; Held *et al*., 2019; Hopke *et al*., 2021). Expected phenotypes involved apical and lateral branching, tracking or nestling, as well as hit-and-split events. In this study, we identified a *reversible* cytoplasmic retraction, which involved withdrawal of cytoplasm, septa formation, breaching of septa and reintroduction of cytoplasm into previously emptied hyphae. Generally, it was observed that hyphae try to grow around an obstacle upon encountering it; they track its edge until it can “escape” the obstacle. If the obstacle proves to be impassable, such as in dead-end corners, the fungus answers by branching off into another direction. As AMF are obligate symbionts (i.e., do not possess saprotrophic capabilities) and their sole focus is to find a suitable host as fast as possible (Smith *et al*., 2010), hyphae and spores have to be highly economical with their resources in the explorative pre-symbiotic state. Therefore, when the original growth path is blocked or appears unfavourable, they retract cytoplasm containing all cellular contents from the hyphal tip in order to redistribute it to another branch of the hyphal network. This retraction is a common observation in filamentous fungi in general (Martínez-Camacho *et al*., 2011), as well as AMF (Kokkoris *et al*., 2020), usually as a defence mechanism to keep the mycelial network from global compromisation (Logi *et al*., 1998; Purin *et al*., 2011). The novelty of our finding, however, is that this process is reversible, as observed in *Rhizophagus* MUCL 41833. The septa, which are formed to segregate empty compartments within a hypha from those that are filled, can be breached and cellular contents reintroduced into the emptied segments. Opening and closing of septa in septate fungi is a well-studied and common process. In Ascomycetes, for example, a so-called Woronin body functions as a plug for septa pores allowing for a controlled, selective permeability through these septa (Bleichrodt *et al*., 2015). In non-septate fungi like AMF, where septation is of a primarily defensive nature, septal opening and closing has not been thoroughly described to date. Lee (2011) proposed the removal of septa to control the expansion of cytoplasmic contents into new hyphal area in *Rhizophagus*. It can be assumed that septa are broken down entirely, as well as being newly formed, when compartmentalisation is needed. The observed reversible cytoplasmic retraction with septation, together with branching, makes hyphal exploration a highly dynamic process. The repopulating of emptied hyphae might be an attempt to re-explore avenues to adapt to a changing environment, again with minimised resource requirements. Besides the cytoplasmic retraction triggered by physical obstacles, it also happens randomly in free hyphae (Bago *et al*., 1998). Which signal or stimulus induces the retraction here is unknown. Observing the dynamic, reversible cytoplasmic retraction was made possible by our high-resolution, real-time microfluidic approach and may have been overlooked by *in vitro* root organ cultures due to the lack of hyphal confinement.

To further characterise the reversible retraction of cytoplasm observed during hyphal space searching, we introduced the fluorescent dyes FM4-64 and calcofluor white into the *AMF-SporeChip* to stain hyphae. While FM4-64 is widely used with filamentous fungi, predominantly for detection of the Spitzenkörper (Fischer-Parton *et al*., 2000; Dijksterhuis *et al*., 2013; Garduño-Rosales *et al*., 2022), calcofluor white is more commonly used in medical studies for the detection of fungal pathogens in tissue samples (Hageage *et al*., 1984; Zhang *et al*., 2010; Chen *et al*., 2021). Both dyes stained the fungal hyphae successfully and confirmed the identity of emptied hyphae. As expected, the chitin-specific calcofluor white stained every hypha, while the lipid bilayer-specific FM4-64 only stained certain hyphae, which usually appeared phase-dark under phase contrast, corroborating the finding that hyphae were emptied of their cytoplasmic content. In these empty hyphal “shells”, only a very faint signal at 544 nm was visible; however, due to a slight autofluorescence of the studied strains it cannot be known with certainty whether the cell membrane remains behind after retraction of the cytoplasm, or rather is removed completely or decays shortly after the event. Implementation of fluorescent dyes such as these, especially FM4-64, for live-cell imaging of AMF hyphae in combination with our microdevices will help to characterise the dynamic retraction process in further detail, as well as aid examination of AMF hyphae for a potential Spitzenkörper.

### Conclusion and outlook

We have presented the first microfluidic device for studying AMF germination, pre-symbiotic hyphal development, anastomosis formation and space searching at the cellular level, which has provided new insights into AMF hyphal growth dynamics and revealed an intricate mechanism of space searching involving reversible cytoplasmic retraction, branching and directional changes. In the future, it is envisaged that the *AMF-SporeChip* could be modified easily to investigate several new frontiers in AMF research. One example includes modifying the design to accommodate two different strains of AMF simultaneously for the study of anastomosis phenotypes in a systematic manner. Further, our device could equally be used as a suitable platform to gain a deeper understanding of septa, as well as hyphopodium formation in AMF. To study metabolites involved in signalling, microfluidics could offer the opportunity to collect the fluidic volume for downstream metabolomics analyses, following visual examinations of these events. Our design is not only limited to AMF, as other fungal spores can be introduced into the *AMF-SporeChip* in a similar manner. Moreover, gradients of chemicals or nutrients can be implemented for studying foraging and signalling in soil exploration. The further development of our device could be also utilised for experiments on AMF interactions with bacteria, e.g., phosphate solubilising bacteria, which have been reported to be important for phosphate uptake in AMF (Jiang *et al*., 2021), suggesting a tripartite symbiosis between plant, fungus and bacteria. To study the AMF symbiosis with their host roots in more detail, a device built from a combination of the *AMF-SporeChip* and the *RootChip* (Stanley *et al*., 2018) could be of assistance.

## Supporting information

Description of Additional Supplementary Files

## Acknowledgments

We acknowledge financial support from the Department of Bioengineering at Imperial College London, as well as the Swiss National Science Foundation in the form of an Ambizione Career Grant (PZ00P2_168005) to C.E.S.

## Author contributions

The idea for the *AMF-SporeChip* and study was conceived by C.E.S. F.R. designed and fabricated the microfluidic device, and performed the experiments, the microscopy imaging and the data analysis. M.C. prepared the spore samples, as well as advice and training with regard to setting up AMF *in vitro* cultures. C.E.S. supervised this study, whereas S.D. and M.H. provided valuable feedback to the experimental work. F.R. and C.E.S. took the lead in writing the manuscript with contributions from all other authors. All authors read and approved the final version of the manuscript.

## Data availability

All relevant data are available from the corresponding author upon request.

## Supplementary information

- **Supplementary Information**
- **Description of Additional Supplementary Files**
- **Supplementary Movie 1**
- **Supplementary Movie 2**
- **Supplementary Movie 3**
- **Supplementary Movie 4**
- **Supplementary Movie 5**
- **Supplementary Movie 6**
- **Supplementary Movie 7**
- **Supplementary Movie 8**
- **Supplementary Movie 9**

