## Supplementary Information for "*AMF-SporeChip* provides new insights into arbuscular mycorrhizal fungal pre-symbiotic hyphal growth dynamics at the cellular level"

### Supplementary figures

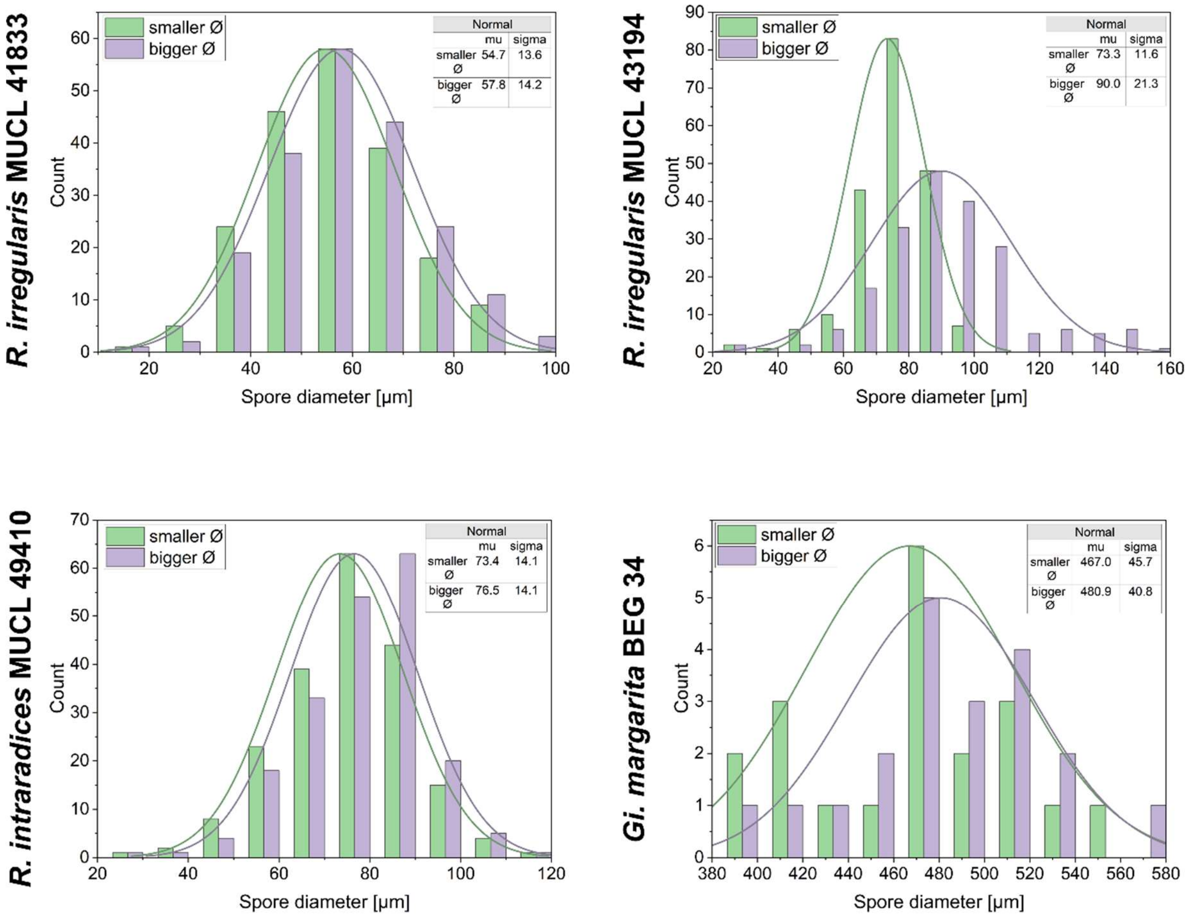


**Supplementary Fig. 1**: Spore diameters of arbuscular mycorrhizal fungi (AMF) used in this study. The diameters of a total of 200 spores per strain (20 spores for *Gigaspora margarita* BEG 34) were measured using the line tool in ImageJ/Fiji and the size distribution plotted with Origin. The largest and smallest diameters of each spore were measured. For *Rhizopagus irregularis MUCL 41833* and *Rhizophagus intraradices MUCL 49410* the difference was very small; for *Rhizopagus irregularis* MUCL 43194, however, a clear difference was observed. Since the total spore number for *Gi. margarita* BEG 34 was very limited, the distribution is quite irregular. For each diameter, the median value “mu (µ)” with a standard deviation “sigma (σ)” is given.

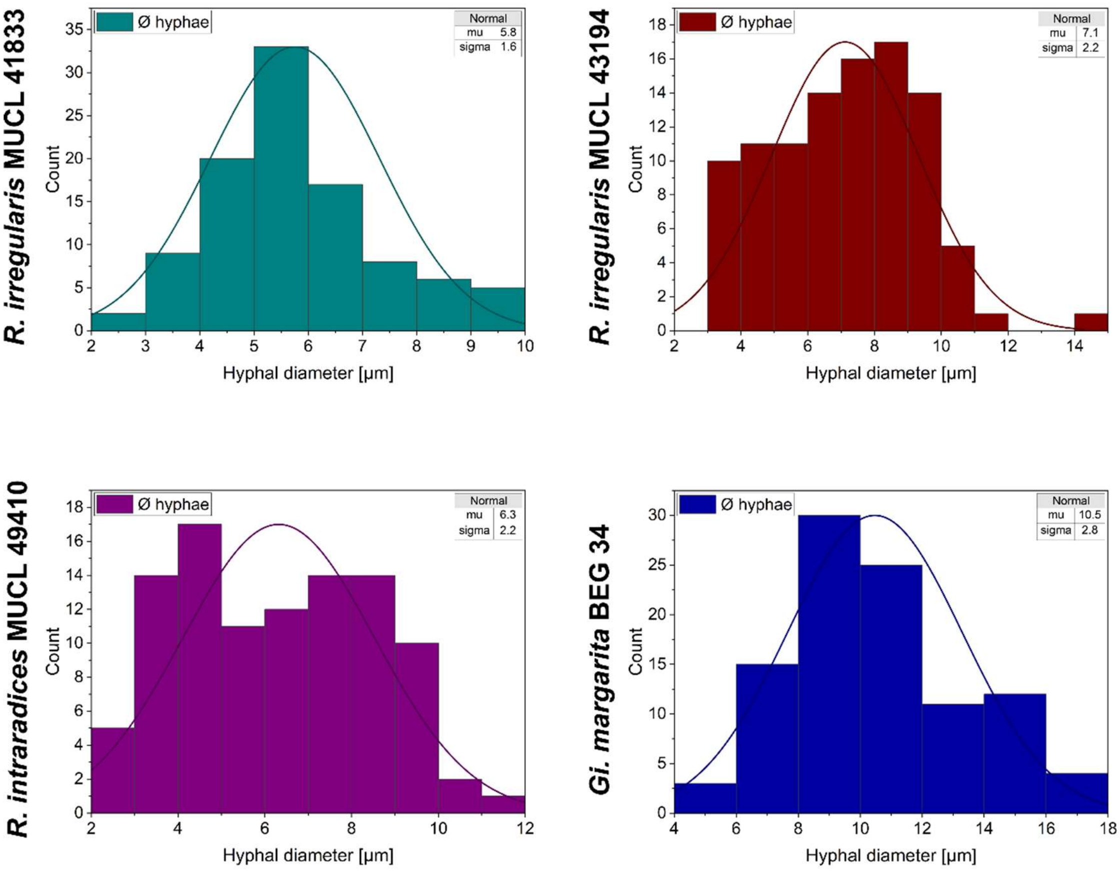


**Supplementary Fig. 2**: Hyphal diameters of arbuscular mycorrhizal (AMF) strains used in this study. The diameters of a total of 100 hyphae per strain were measured in random spots on the hypha using the line tool in ImageJ/Fiji and the size distribution plotted with Origin. For each diameter, the median value "mu (µ)" with a standard deviation "sigma (σ)" is given.

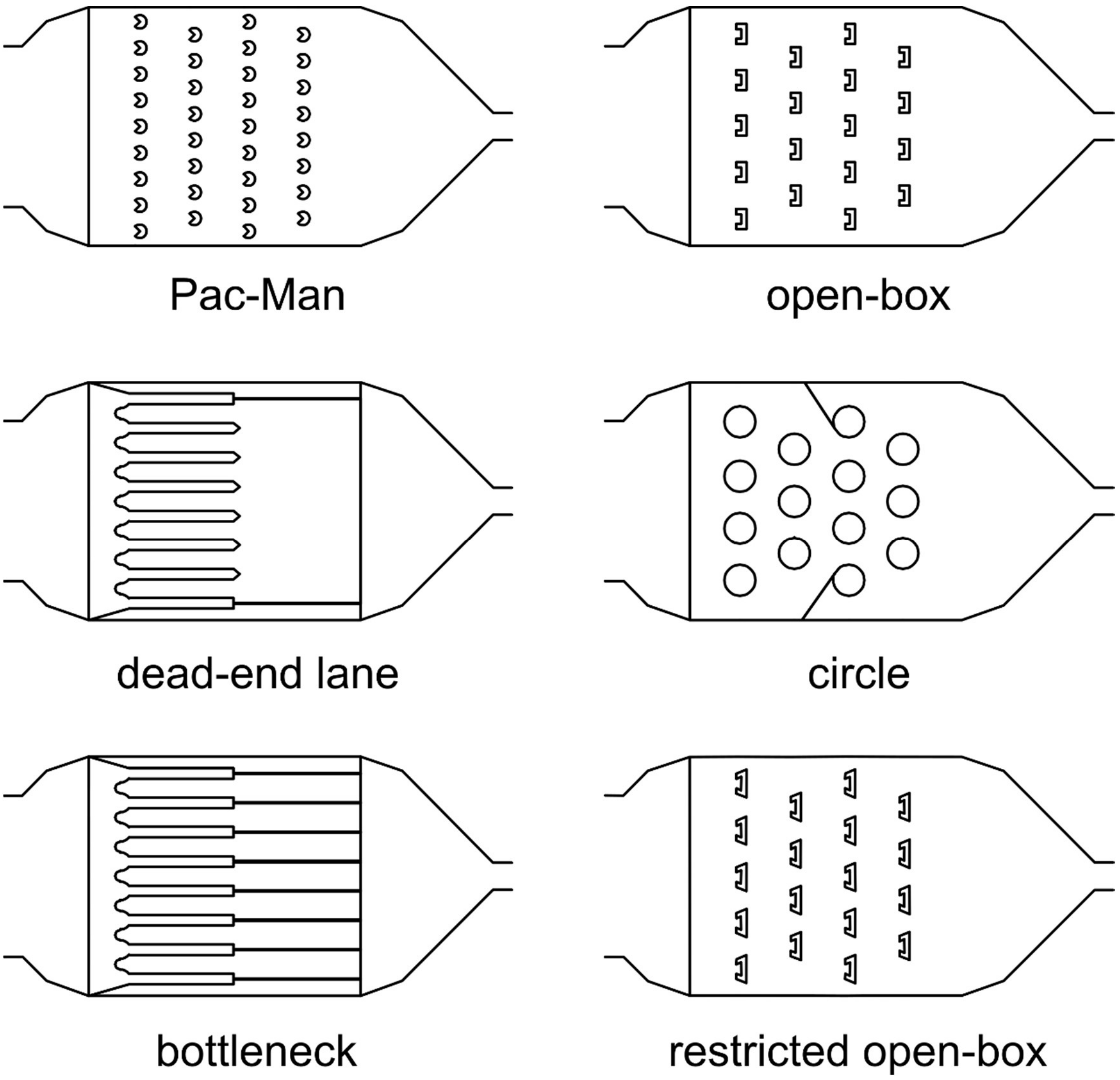


**Supplementary Fig. 3**: Different physical obstacles accommodated in the investigation zone of the *AMF-SporeChip*.

### Supplementary methods

#### *Media preparation*

For the preparation of MSR(-) medium, used for the hyphal compartment of the arbuscular mycorrhizal fungi (AMF) co-culture, 0.59 g Modified Strullu and Romand (MSR, Duchefa Biochemie, the Netherlands) powder was dissolved in 1 l sterile double distilled water (ddH2O) containing 10 ml of a 0.15 M calcium nitrate solution. The pH was adjusted to 5.5 using 0.1 M NaOH and HCl, and finally 3 g phytagel (Merck, Germany) added. MSR medium, used for chicory root culture and the root compartment of the AMF co-culture, was prepared in the same manner, however, 10 g sucrose were added to the solution before adjusting the pH (Declerck *et al.*, 1998; Cranenbrouck *et al.*, 2005). After autoclaving, the media must be kept at 60 °C to prevent it from solidifying. Once cooled down and solidified it cannot be melted again. For the on-chip experiments, a liquid version of the MSR(-) medium was prepared; the pH was adjusted to 7 and no phytagel was added (the pH of the phytagel-containing media was measured to be 7 before autoclaving).

#### *AMF culture*

The AMF strains *Rhizophagus irregularis* MUCL 43194, *Rhizophagus irregularis* MUCL 41833 and *Rhizophagus intraradices* MUCL 49410 were maintained *in vitro* in bi-compartmented Petri plates (Ø = 90 mm) on Ri T-DNA transformed roots of chicory (*Cichorium intybus* L.) on MSR medium. One compartment was filled with 15-20 ml of MSR(-) (termed the "hyphal compartment"), such that the liquid medium is held by the central wall due to surface tension. Once the MSR(-) medium has solidified, 5-15 ml of MSR medium was carefully poured into the second compartment (termed the "root compartment") while tilting the plate towards the hyphal compartment (i.e., to ensure connectivity between the two compartments). The root compartment was then inoculated with a fresh and vital chicory root, the plates were then sealed with cling film and incubated at 27 °C in the dark in inverted position for one week. Plugs of medium containing spores from an at least 3-month-old culture plate showing a high spore density were selected and transferred to an empty sterile Petri plate. These plugs were then cut into very thin pieces and mixed with a scalpel. The viscous spore-containing paste was then spread directly onto the chicory roots. The plates were then sealed with cling film and incubated at 27 °C in the dark in inverted position until the spores were mature (usually after 3-6 months).

Spores of *Gigaspora margarita* BEG 34 were produced in pot culture in double-autoclaved (121°C for 15 min) substrate composed of thin quartz (0.4-0.8 mm), larger quartz (1-2 mm) and calcinated clay (2/1/2; v/v/v) with *Plantago lanceolata*. After at least 9 months of cultivation, spores were sampled and disinfected as described by Cranenbrouck *et al.* (2005). Briefly, spores were disinfected using a filtration apparatus connected to a vacuum outlet. First, they were cleaned twice in sterile water, then treated with chloramine T 2% solution (with 2 drops of Tween 20) for 10 min. Spores were then washed again 3 times with sterile water before treatment with a filtered (using a 0.22 µm Acrodisk filter) antibiotic solution composed of streptomycine sulfate 0.02% and gentamycine sulfate 0.01%, for 10 min. Spores were stored in sterile water at 4°C before use.

***Microfluidic device fabrication***

The device designs were drawn in AutoCAD 2022 (Autodesk) and checked for correct structuring using KLayout (Köfferlein, 2006). The design was then printed to create a mylar® film photolithography mask by Micro Lithography Services Ltd., UK. For the master mould manufacturing (conducted in a clean room), a fresh silicon wafer (Diameter: 4"; Orientation: <100>; Dopant: P(Boron); Resistivity: 0-100 ohm-cmCz; Centre Thickness: 425-550 µm; Surface: Single side polished, Si-Mat Silicon Materials, Germany) was washed with acetone and isopropanol and subsequently plasma-cleaned for 5 min using a plasma cleaner (PDC-002-CE, Harrick Plasma, USA). The following procedure was used to prepare the master mould for *Rhizophagus* spp. devices, with parameters for the *Gi margarita* master mould written in parentheses if deviating from the standard protocol. For the first layer, 5 ml of SU-8 2010 (Kayaku, USA) were spin-coated onto the wafer, spinning at 500 rpm for 10 s and 3700 rpm for 30 s (Spin Coater WS-650MZ Modular, Laurell, USA), followed by soft baking at 65 °C for 30 s and 95 °C for 2.5 min ramping up with 5 °C/min on a hot plate (Hot Plate EMS 1000-3, Ascon Tecnologic, Italy). In the next step, the wafer was exposed to ultraviolet (UV) light at a dose of 130 mJ/cm$^2$ using a UV KUB-3 mask aligner (Kloé SA, France) and baked at 65 °C for 30s and 95 °C for 3.5 min, ramping up with 5 °C/min. For the second layer, 5 ml SU-8 2075 (Kayaku, USA) (SU-8 2150 (Kayaku, USA)) were spun onto the first layer at 500 rpm for 20 s and

2200 rpm (1300 rpm) for 30 s, followed by soft baking at 65 °C for 5 min (7 min) and 95 °C for 15 min (60 min) ramping up with 5 °C/min on a hot plate. For a better alignment of the first and second layer, the SU-8 2010 was removed from the alignment marks using acetone and clean room sampling swab (Berkshire, UK). The layers were aligned and the wafer exposed with 228 mJ/cm$^2$ (450 mJ/cm$^2$) UV light using the UV KUB-3 mask aligner and subsequently baked at 65 °C for 3.5 min (5 min) and 95 °C for 9 min (35 min) ramping up with 5 °C/min on a hot plate. For the development step, the wafer was agitated in SU-8 developer (2-(1-methoxy)propyl acetate, 99%, Thermo Scientific, USA) for 8 min (16 min), then for 1 min (4 min) with fresh developer and finally washed with isopropanol and dried with an airgun. In the last step, the master mould was hard baked at 150 °C for 2 min (5 min) on a hot plate.

***Imaging***

To obtain time-lapse images (used to produce growth and germination videos) as well as time-point large images (used for measuring growth rates) two inverted microscopes (Eclipse Ti-U and Eclipse Ti-2, Nikon) were employed. The microscopes were equipped with air immersed ×10/0.3 NA (numerical aperture) and x20/0.45 NA Plan Fluor objectives (Nikon; used for time-point images and time-lapse experiments respectively), motorised stages and camera heads (Ti-U: Retiga R1 CCD camera (Qimaging, Canada); Ti-2: DS-Qi2 Mono Digital Microscope Camera (Nikon, UK)). For growth rate experiments, the large image function in NIS-Elements Advanced Research imaging software (Nikon) was used to acquire images once every 24 h for 7 days, with a final image being taken after 14 days. To cover the entire device, 9 × 5 fields of view were needed; the acquired image array was then automatically stitched together with a 20% overlap. For long term microscopy, a temperature-controlled incubator (Okolab, Italy) mounted around the microscope stage was set to 27 °C.

The Eclipse Ti-U was used for fluorescence imaging only, being equipped with a high-power light emitting diode (LED) light engine (LedHUB, Omicron-Laserage Laserprodukte GmbH, Germany). Two LEDs, with peak wavelengths of ca. 385 nm and ca. 505-600 nm, were employed to excite AMF strains dyed with calcofluor white and FM4-64 respectively. The combination of dyes was chosen for their tissue specificity as well as good spectral separation, with calcofluor white and FM4-64 exhibiting emission maxima of 432 nm and 670 nm, respectively (Au - Lichius *et al.*, 2019). In this study, the dyes were used solely for end-point staining. Hence, any potential toxic effects could be neglected, i.e., associated with the use of calcofluor white, which has been reported to disrupt normal formation of chitin microfibrils due to its intercalating nature (Herth, 1980).

The following filters were used: (i) DAPI filter set (AHF Analysentechnik AG, Germany) with Excitation filter: 377/50 BrightLine HC (352-403 nm), Beamsplitter: 409 nm and Emission filter: 447/60 BrightLine HC (417-477 nm) and (ii) TRITC filter set with Bandpass filter: 495 nm (Omicron-Laserage Laserprodukte GmbH, Germany), Beamsplitter: 565 nm (AHF Analysentechnik AG, Germany) and Emission filter: 605/70 nm (570-640 nm) (AHF Analysentechnik AG, Germany).

***On-plate experiment***

For the on-plate experiment, spores were picked up with a pipette (Pipetman G, 200 µl, Gilson, USA) and evenly distributed on a glass-bottomed Petri dish (Ø dish = 50 mm, Ø glass = 40 mm; Fluorodish, World Precision Instruments, Germany) containing a thin layer of MSR(-) (ca. 1-2 mm). The plates were then sealed with cling film and incubated at 27 °C. Spore germination and hyphal growth were imaged and measured alongside the microdevices in the same manner.
