## Supplementary material for "*AMF-SporeChip* provides new insights into arbuscular mycorrhizal fungal pre-symbiotic hyphal growth dynamics at the cellular level": Description of Additional Supplementary Files

**File name: Supplementary Movie 1**

**Description: Germination of *Rhizophagus irregularis* MUCL 41833 within the *AMF-SporeChip*.** A time-lapse experiment was recorded over 19 hours and 15 minutes with 15 min intervals between image acquisitions using phase contrast microscopy. The movie shows the germination of one spore of *Rhizophagus irregularis* MUCL 41833 in a microchannel with a single, straight hypha emerging from the germination site. Scale bar = 100 µm. Time stamp format = hh:mm:ss.

**File name: Supplementary Movie 2**

**Description: Germination of *Rhizophagus irregularis* MUCL 43194 within the *AMF-SporeChip*.** A time-lapse experiment was recorded over 55 hours with 20 min intervals between image acquisitions using phase contrast microscopy. The movie shows the germination of several spores of *Rhizophagus irregularis* MUCL 43194 in a microchannel with multiple, curly hyphae emerging from each germination site. In some spores, prior to germination, a change in phase contrast down the subtending hypha can be observed, indicating a directed discharge of cellular contents. Scale bar = 50 µm. Time stamp format = hh:mm:ss.

**File name: Supplementary Movie 3**

**Description: Germination of *Rhizophagus irregularis* MUCL 49410 within the *AMF-SporeChip*.** A time-lapse experiment was recorded over 78 hours with 20 min intervals between image acquisitions using phase contrast microscopy. The movie shows the germination of one spore of *Rhizophagus irregularis* MUCL 49410 in a microchannel with a single hypha emerging from the germination site, which branches several times, shortly behind the initial point of germination. Scale bar = 100 µm. Time stamp format = hh:mm:ss.

**File name: Supplementary Movie 4**

**Description: Spores of *Rhizophagus irregularis* MUCL 49410 internally collapsing within the *AMF-SporeChip*.** A time-lapse experiment was recorded over 23 hours and 20 minutes with 20 min intervals between image acquisitions using phase contrast microscopy. The movie shows spores of *Rhizophagus irregularis* MUCL 49410 collapsing internally, and suddenly ejecting storage vesicles and cellular content into the attached hyphae. Scale bar = 100 µm. Time stamp format = hh:mm:ss.

**File name: Supplementary Movie 5**

**Description: Hypha of *Gigaspora margarita BEG 34* growing into the *AMF-SporeChip*.** A time-lapse experiment was recorded over 25 hours and 20 minutes with 20 min intervals between image acquisitions using phase contrast microscopy. The movie shows a hypha of *Gigaspora margarita* BEG 34 growing into a dead-end microchannel. Hyphal branching, cytoplasmic retraction and septa formation can be observed. Scale bar = 150 µm. Time stamp format = hh:mm:ss.

**File name: Supplementary Movie 6**

**Description: Tip-to-tip anastomosis between two hyphae of *Rhizophagus irregularis* MUCL 43194 within the *AMF-SporeChip*.** A time-lapse experiment was recorded over 21 hours with 20 min intervals between image acquisitions using phase contrast microscopy. The movie shows two hyphae of *Rhizophagus irregularis* MUCL 43194 approaching each other in a directed, stop-and-go manner, prior to tip-to-tip anastomosis. Scale bar = 100 µm. Time stamp format = hh:mm:ss.

**File name: Supplementary Movie 7**

**Description: Tip-to-tip anastomosis between a hypha and a germination site of *Rhizophagus irregularis* MUCL 43194 within the *AMF-SporeChip*.** A time-lapse experiment was recorded over 83 hours and 40 minutes with 20 min intervals between image acquisitions using phase contrast microscopy. The movie shows a hypha of *Rhizophagus irregularis* MUCL 43194 approaching a germination site of a subtending hypha, to which it anastomoses. After 57 h, a change in phase contrast shifting from the subtending hypha into the approaching hypha can be observed, suggesting a transfer of cellular contents through the newly formed connection, followed by the formation of new branches. Scale bar = 50 µm. Time stamp format = hh:mm:ss.

**File name: Supplementary Movie 8**

**Description: Tip-to-side anastomosis between two hyphae of *Rhizophagus irregularis* MUCL 43194 within the *AMF-SporeChip*.** A time-lapse experiment was recorded over 20 hours and 20 minutes with 20 min intervals between image acquisitions using phase contrast microscopy. The movie shows a hypha of *Rhizophagus irregularis* MUCL 43194 approaching a second hypha and anastomoses with it from the side. After successful merging, fluctuation of cellular contents between both hyphae can be observed. Scale bar = 25 µm. Time stamp format = hh:mm:ss.

**File name: Supplementary Movie 9**

**Description: Dynamic hyphal reactions of *Rhizophagus irregularis* MUCL 41833 within the *AMF-SporeChip*.** A time-lapse experiment was recorded over 45 hours and 45 minutes with 15 min intervals between image acquisitions using phase microscopy. The movie shows the growth of a hypha of *Rhizophagus irregularis* MUCL 41833 being obstructed by an obstacle, causing a change in growth behaviour, involving growth arrest, cytoplasmic retraction, directional changes, lateral branching as well as reversal of cytoplasmic retraction. Scale bar = 100 µm. Time stamp format = hh:mm:ss.
